# deadtrees.earth - An Open-Access and Interactive Database for Centimeter-Scale Aerial Imagery to Uncover Global Tree Mortality Dynamics

**DOI:** 10.1101/2024.10.18.619094

**Authors:** Clemens Mosig, Janusch Vajna-Jehle, Miguel D. Mahecha, Yan Cheng, Henrik Hartmann, David Montero, Samuli Junttila, Stéphanie Horion, Mirela Beloiu Schwenke, Stephen Adu-Bredu, Djamil Al-Halbouni, Matthew Allen, Jan Altman, Claudia Angiolini, Rasmus Astrup, Caterina Barrasso, Harm Bartholomeus, Benjamin Brede, Allan Buras, Erik Carrieri, Gherardo Chirici, Myriam Cloutier, KC Cushman, James W. Dalling, Jan Dempewolf, Martin Denter, Simon Ecke, Jana Eichel, Anette Eltner, Maximilian Fabi, Fabian Fassnacht, Matheus Pinheiro Feirreira, Julian Frey, Annett Frick, Selina Ganz, Matteo Garbarino, Milton García, Matthias Gassilloud, Marziye Ghasemi, Francesca Giannetti, Roy Gonzalez, Carl Gosper, Konrad Greinwald, Stuart Grieve, Jesus Aguirre Gutierrez, Anna Göritz, Peter Hajek, David Hedding, Jan Hempel, Melvin Hernández, Marco Heurich, Eija Honkavaara, Tommaso Jucker, Jesse M. Kalwij, Pratima Khatri-Chhetri, Hans-Joachim Klemmt, Niko Koivumäki, Kirill Korznikov, Stefan Kruse, Robert Krüger, Etienne Laliberté, Liam Langan, Hooman Latifi, Jan Lehmann, Linyuan Li, Emily Lines, Javier Lopatin, Arko Lucieer, Marvin Ludwig, Antonia Ludwig, Päivi Lyytikäinen-Saarenmaa, Qin Ma, Giovanni Marino, Michael Maroschek, Fabio Meloni, Annette Menzel, Hanna Meyer, Mojdeh Miraki, Daniel Moreno-Fernández, Helene C. Muller-Landau, Mirko Mälicke, Jakobus Möhring, Jana Müllerova, Paul Neumeier, Roope Näsi, Lars Oppgenoorth, Melanie Palmer, Thomas Paul, Alastair Potts, Suzanne Prober, Stefano Puliti, Oscar Pérez-Priego, Chris Reudenbach, Christian Rossi, Nadine Katrin Ruehr, Paloma Ruiz-Benito, Christian Mestre Runge, Michael Scherer-Lorenzen, Felix Schiefer, Jacob Schladebach, Marie-Therese Schmehl, Selina Schwarz, Rupert Seidl, Elham Shafeian, Leopoldo de Simone, Hormoz Sohrabi, Laura Sotomayor, Ben Sparrow, Benjamin S.C. Steer, Matt Stenson, Benjamin Stöckigt, Yanjun Su, Juha Suomalainen, Michele Torresani, Josefine Umlauft, Nicolás Vargas-Ramírez, Michele Volpi, Vicente Vásquez, Ben Weinstein, Tagle Casapia Ximena, Katherine Zdunic, Katarzyna Zielewska-Büttner, Raquel Alves de Oliveira, Liz van Wagtendonk, Vincent von Dosky, Teja Kattenborn

## Abstract

Excessive tree mortality is a global concern and remains poorly understood as it is a complex phenomenon. We lack global and temporally continuous coverage on tree mortality data. Ground-based observations on tree mortality, *e.g*., derived from national inventories, are very sparse, not standardized and not spatially explicit. Earth observation data, combined with supervised machine learning, offer a promising approach to map tree mortality over time. However, global-scale machine learning requires broad training data covering a wide range of environmental settings and forest types. Drones provide a cost-effective source of training data by capturing high-resolution orthophotos of tree mortality events at sub-centimeter resolution. Here, we introduce deadtrees.earth, an open-access platform hosting more than a thousand centimeter-resolution orthophotos, covering already more than 300,000 ha, of which more than 58,000 ha are fully annotated. This community-sourced and rigorously curated dataset shall serve as a foundation for a global initiative to gather comprehensive reference data. In concert with Earth observation data and machine learning it will serve to uncover tree mortality patterns from local to global scales. This will provide the foundation to attribute tree mortality patterns to environmental changes or project tree mortality dynamics to the future. Thus, the open and interactive nature of deadtrees.earth together with the collective effort of the community is meant to continuously increase our capacity to uncover and understand tree mortality patterns.

## 1 Introduction

In recent decades, elevated tree mortality rates have been reported for many regions of the world (Hartmann et al. 2022). This phenomenon is attributed to climate change-induced more frequent and intense climate extremes such as droughts, heatwaves, and late frosts, that often trigger outbreaks of damaging insects or epidemic diseases (Anderegg et al. 2013; Bauman et al. 2022; Gora and Esquivel-Muelbert 2021; Hartmann et al. 2022; Senf et al. 2020; Trumbore et al. 2015). Tree mortality is generally not driven by a single driver but by complex compound events, consisting of multiple biotic and abiotic agents and feedbacks (Allen et al. 2010; Bastos et al. 2023; Mahecha et al. 2024). This may include a combination of consecutive heatwaves, meteorological and soil droughts, followed by late frosts after leaf budding, and the infestation of already weakened trees by pest and pathogens (Coleman et al. 2018; Fettig et al. 2019; Stephenson et al. 2019; Trugman et al. 2021).

Trees are long-lived and sessile organisms that cannot escape extreme conditions via migration, and their capacity to acclimate or adapt evolutionary to rapid environmental changes is slow (Allen et al. 2015). Accordingly, the spatio-temporal patterns of standing dead tree canopies are direct indicators of how different tree species, functional types, ages, or entire ecosystems cope with biotic and abiotic stressors (Anderegg et al. 2013; Hartmann et al. 2022). Moreover, timely information on tree mortality dynamics is urgently needed by decision-makers in forest management and nature conservation. Information on tree mortality patterns is required to identify adaptation strategies, including selecting tree species, optimizing harvesting cycle, managing pest and disease outbreaks (*e.g*., bark beetle), ensure the provision of ecosystem services and controlling fuel accumulation for wildfire risk reduction (Garrity et al. 2013; Moghaddas et al. 2018; Stephens et al. 2018, 2022; Vilanova et al. 2023; Winter et al. 2024). Moreover, tracking tree mortality patterns helps indicate where ecosystems are undergoing rapid compositional transformations, *i.e*., shift in species and their role in the terrestrial carbon cycle, *e.g*., via declining net carbon sinks (Hill et al. 2023; Pan et al. 2011; Scheffer et al. 2001; Stephens et al. 2022).

Despite its importance, the extent and rate of tree mortality at the global scale remains largely unknown or imprecise (Allen et al. 2015). Although ground-based inventories are the gold standard in forestry, national forest inventories only sometimes record tree mortality, but usually have sparse spatial coverage (Puletti et al. 2019) and low temporal sampling frequencies (*e.g*., 10-year intervals), which do not align well with the rapid dynamics of environmental stressors. Therefore, these inventories provide limited assistance in attributing tree mortality to short-term environmental dynamics such as climate extremes or insect outbreaks (Hülsmann et al. 2017; Woodall et al. 2005). Consequently, meta-analyses based on such ground observations could be biased or underrepresented for recent elevated tree mortality (Hammond et al. 2022; Yan et al. 2024). The value of field inventories for global tree mortality studies is further complicated by the commonly low data accessibility and heterogeneity in sampling protocols and data quality (McRoberts et al. 2010; Senf et al. 2018). Recent initiatives such as the global tree mortality database (Hammond et al. 2022) have gathered and har-monized invaluable information towards a global assessment of tree mortality. However, they are still severely limited in their spatial and temporal coverage and are not based on a systematic assessment that would enable scaling to larger spatial scales. Uncovering global tree mortality patterns requires a multi-faceted approach that complements the ground-based assessments.

Satellite-based Earth Observation offers a promising avenue, providing seamless spatial coverage and temporally consistent monitoring (The International Tree Mortality Network et al. 2024). Using data from the Landsat satellite mission, Hansen *et al*. created the prominent global *forest loss* map by applying a decision tree classifier on time series of spectral metrics (Hansen et al. 2013). However, this approach reveals a binary classification of forest loss, not tree mortality, and is restricted to 30 m spatial resolution and thus cannot detect the often scattered patterns of tree mortality (Cheng et al. 2024; Espírito-Santo et al. 2014; Schiefer et al. 2024). Unsupervised approaches, that is analysis without labeled reference data, can reveal continuous forest responses using anomalies of vegetation indices, which are computed by combining multiple spectral bands for each pixel (Lange et al. 2024; Senf and Seidl 2021; Senf et al. 2018, 2020; Thonfeld et al. 2022). However, vegetation indices cannot directly reveal tree mortality and using such methods to uncover scattered and small-scale mortality remains a challenging task. Therefore, translating the complex Earth observation signals to tree mortality patterns requires a supervised approaches (Schiefer et al. 2023).

The Earth observation community, thus, currently lacks a representative collection of reference data for training and validating supervised methods for monitoring tree mortality. Given the relatively coarse resolution, satellite data does not provide the necessary spatial detail to extract such reference data directly. Airplane aerial images typically have higher resolutions and are often freely available for regions or entire countries and, therefore, provide a promising source to map tree mortality (Cheng et al. 2024; Junttila et al. 2024; Schwarz et al. 2024). However, airplane imagery are only openly available in few countries and their spatial resolutions typically range from 20-60 cm, in rare cases up to 10 cm. This can be a critical constrain to uncover tree mortality, as an image resolution of 20 cm or less does not always enable most precise differentiation of dead from alive tree crowns and may lead to missing small dead trees (compare Figure 1). For some species, crown shapes, or sizes, mortality is still clearly visible at 60 cm and in studies that are limited to specific ecosystems, *e.g*., with dominantly coniferous species, coarse aerial images suffice (Junttila et al. 2024). Such resolution does not suffice, to accurately reveal partial dieback of broadleaf trees (row *B* in Figure 1), mortality atop a bright forest floor (row *C* in Figure 1), or in the presence of objects such as rocks that have a geometry that is similar to tree crowns (*e.g*., rocks, row *D* in Figure 1). Hence, to achieve accurate reference data across all ecosystems and tree types a finer resolution in the centimeter range (*≤* 10 *cm*) is needed, calling for a representative global collection of centimeter-scale imagery.

**Figure 1.**
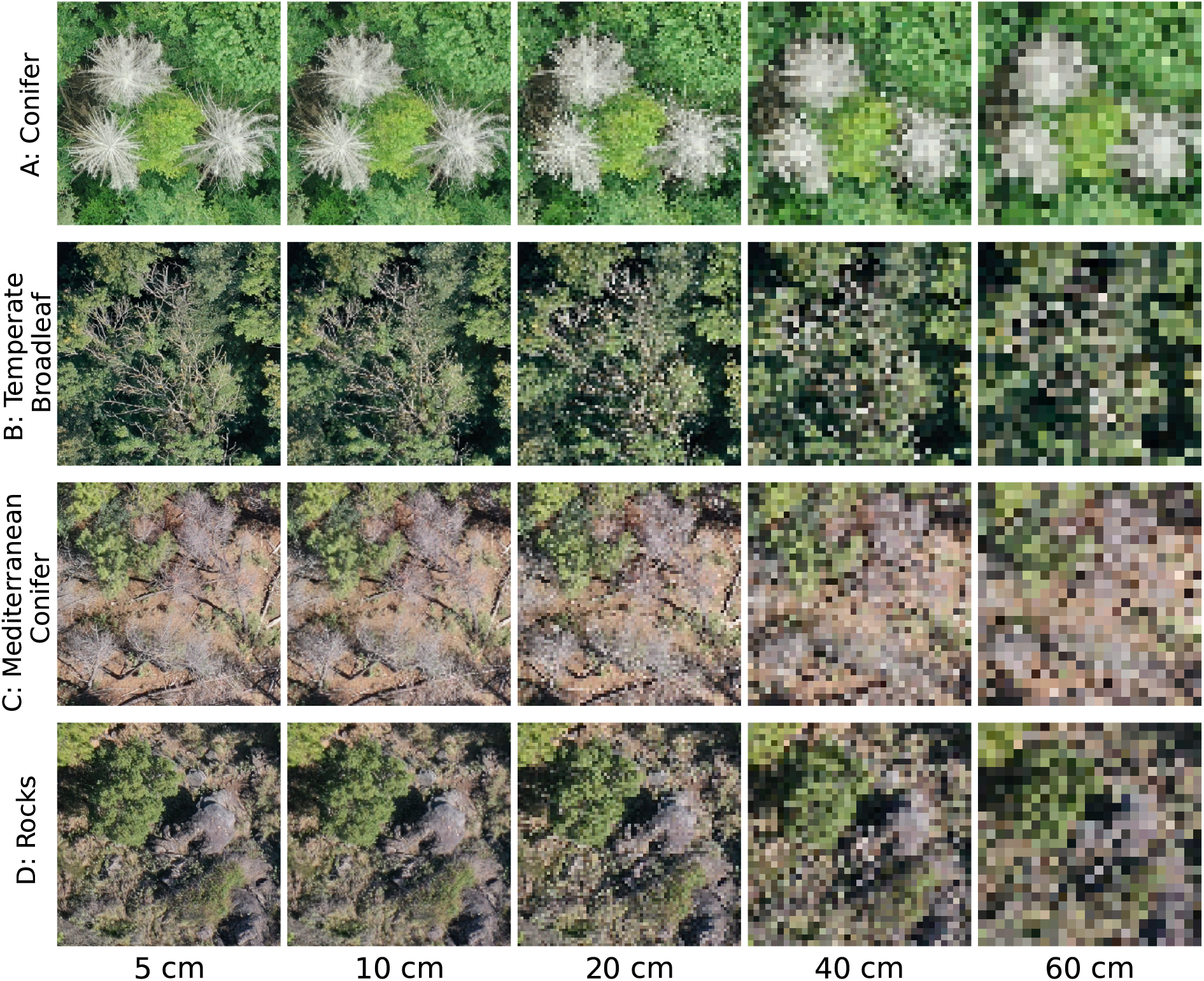
Four forest sites, 15 m in width and height and at resolutions of 5 cm to 60 cm. From top to bottom (*A* to *C*), the tree species are *Picea abies, Fraxinus excelsior*, and *Pinus sylvestris*. Row *D* shows an example where rocks cannot be distinguished from deadwood in coarse-resolution images. The original images have resolutions better than 5 cm and were resampled (nearest-neighbor) for this visualization. Airplane images at the same resolution commonly appear less clear at similar resolutions, hence these images are best-case scenarios.

Drones are becoming increasingly accessible and require minimal training for operation (P. Johnson et al. 2017; Rossi and Wiesmann 2024; Tang and Shao 2015). Suitable orthophotos for precise tree mortality identification at the centimeter scale can be obtained by non-technical users with consumer-type drones and easy-to-use mapping apps. In a recent case study in Germany, Schiefer et al. (2023) leveraged high-resolution drone aerial images (4 cm resolution) as reference to infer the fractional cover of standing deadwood [%] in pixels of satellite data (Sentinel-1 and -2). However, drones require operators to go into the field, creating significant labor costs and time investment. Hence, leveraging drone orthophotos for use in global tree mortality monitoring can only be achieved through a large collective effort across institutions, researchers, and citizens across the globe, to finally acquire a rich collection of orthophotos to represent all forest ecosystems.

Here, we introduce deadtrees.earth, an open science, collaborative platform for accessing, sharing, analyzing, and visualizing a global database of orthophotos with labeled standing deadwood. The deadtrees.earth platform features open-access interactive functionality, allowing users to upload and download images and labels through the website and an API. It also incorporates expert quality control workflows to maintain high data standards. This collection, across spatial and temporal scales, offers unparalleled opportunities for researchers to advance satellite-based model training and validation. The platform’s backend is built with a scalable architecture to allow growth into a large machine learning model ecosystem. Beyond machine-learning applications, this database also enables verification of existing products. Contributors are acknowledged for their data contributions, fostering transparent community participation and acknowledgment.

## 2 The deadtrees.earth platform

deadtrees.earth is a dynamic, community-built, open-access database for aerial orthophotos of delineated standing deadwood. This section presents our definition of standing deadwood, the database structure, database statistics, and a web platform for the integration of the database into the community.

### 2.1 Standing Deadwood

We focus on *standing deadwood*, defined as woody material (twigs, branches, or stems) that has died off but has largely retained its original structure, including brown-stage mortality. For deciduous tree that is a lack of leafs in leaf-on season, that is either in summer or in wet season (Figure 2). Standing deadwood can be identified in centimeter-scale RGB images acquired by drones or airplanes by methods such as semantic segmentation, which involves the generic segmentation of any dead tree crown or branch (Schiefer et al. 2023), or instance segmentation, where each segment corresponds to an individual tree crown (Cheng et al. 2024).

**Figure 2.**
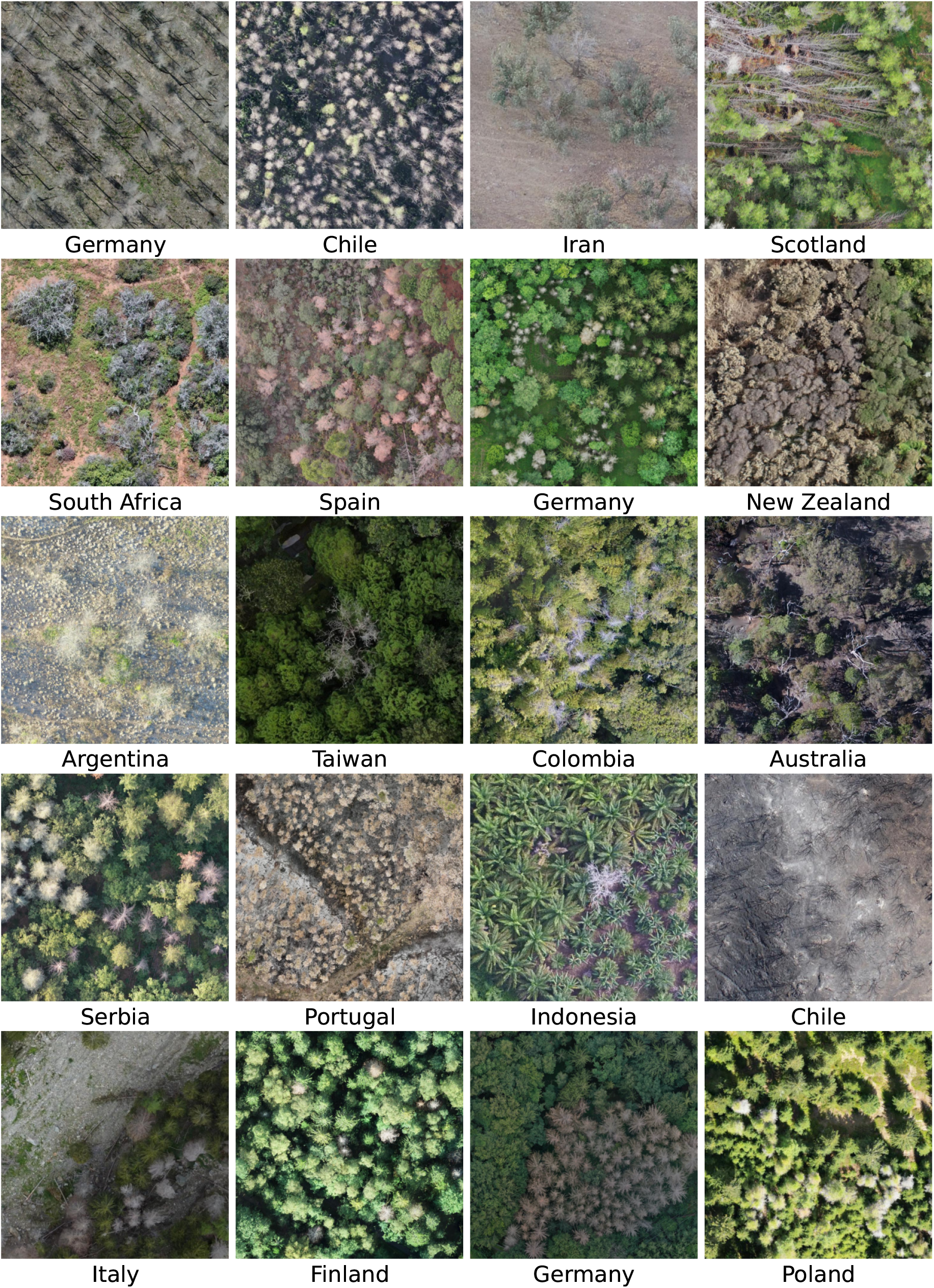
Sample image sections of standing and lying deadwood in a variety of contexts. The caption below each image denotes the acquisition location of the drone orthophoto. All images are available in the database.

Information on lying deadwood is not considered for this database. In contrast to standing dead tree crowns, fallen tree stems are less likely to be detected in drone and airplane imagery, as they are readily occluded by surrounding tree crowns or are rapidly covered by understory. Additionally, fallen trees can be several decades old and are hence less interesting for studying tree mortality as a response to recent environmental changes, climate extremes, or pests and pathogens.

The amount of standing deadwood changes over time with different events (Figure 3). Climate extreme events, such as droughts, can cause tree mortality, increasing the amount of standing deadwood. Standing deadwood is not limited to fully dying trees; partial dieback also affects the amount of standing deadwood. Explicitly including partial dieback is important, as it can be difficult to visually separate trees in imagery of dense forests with complex crown structures (South Africa, Iran, and Australia in Figure 2). In subsequent years, standing dead trees decompose and the fraction of standing deadwood decreases. As soon as dead trees fall over, are felled, or are completely removed, they no longer count as standing deadwood.

**Figure 3.**
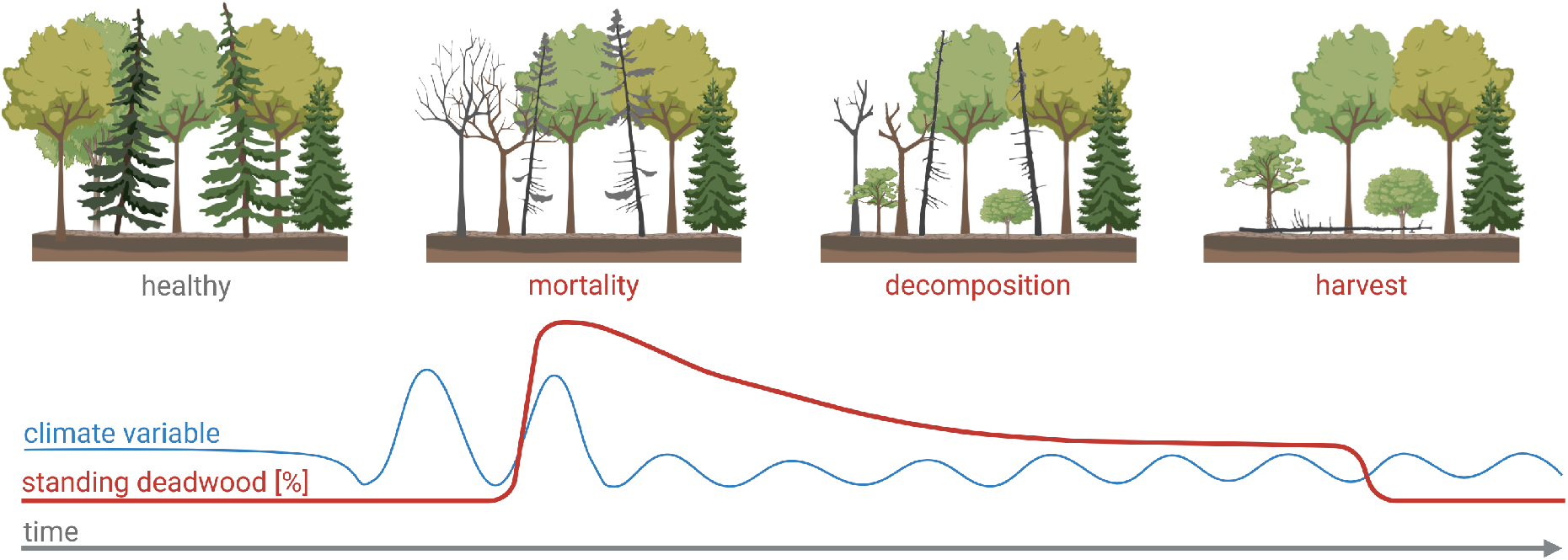
Temporal signature of standing deadwood (red) in multiple scenarios. Climate extreme events (blue) cause tree mortality to increase. Natural decomposition and/or harvesting/salvaging decreases standing deadwood.

Although the concept of standing deadwood is simple, understanding its temporal dynamics requires several considerations. First, the falling of healthy trees does not affect the fraction of standing deadwood. This also includes removing unhealthy trees that have not yet changed their appearance from above and are removed before visible leaf loss. Secondly, a high amount of standing deadwood in one year does not imply that those trees died that year, but several years before that is also possible. Note that the year of the first appearance can be extracted from a standing deadwood time series (Schiefer et al. 2024). Thirdly, drought or cold semi-deciduous species that shed their leaves during climate extremes or species that resprout epicormically after disturbances such as fire, may visually appear as standing deadwood at one time point but may regrow leaves at a later time, *e.g*., red needle cast (Watt et al. 2024).

### 2.2 Database Structure

The deadtrees.earth database is a collection of geo-referenced RGB orthophotos gathered over forests with optionally one or more sets of labels depicting standing deadwood. Our database focuses on airborne imagery better than 10 cm while also allowing submissions of up to 1 m for unrepresented regions or where validated tree mortality labels are provided.

Each **orthophoto** comes with the following metadata: acquisition date, author(s), resolution, platform, resolution and license (compare Figure 4). The author(s) can be one or multiple individuals who contributed to capturing the orthophoto. The acquisition date is crucial for linking with environmental conditions to validate whether the orthophoto was captured in leaf-on season because one cannot differentiate between dead and alive trees in orthophotos that were captured in leaf-off season. Given that data contributors track the acquisition date with different accuracy, we accommodate three levels of precision for the acquisition date, that is, accurate in days, months, or years. Noting the possible temporal error is of utmost importance when combining these observations with other datasets, such as satellite time series (see Subsection 3.2). Also, for each orthophoto, the average ground sampling distance (GSD) is automatically calculated to allow users to filter data based on different spatial resolutions (see Figure 4).

**Figure 4.**
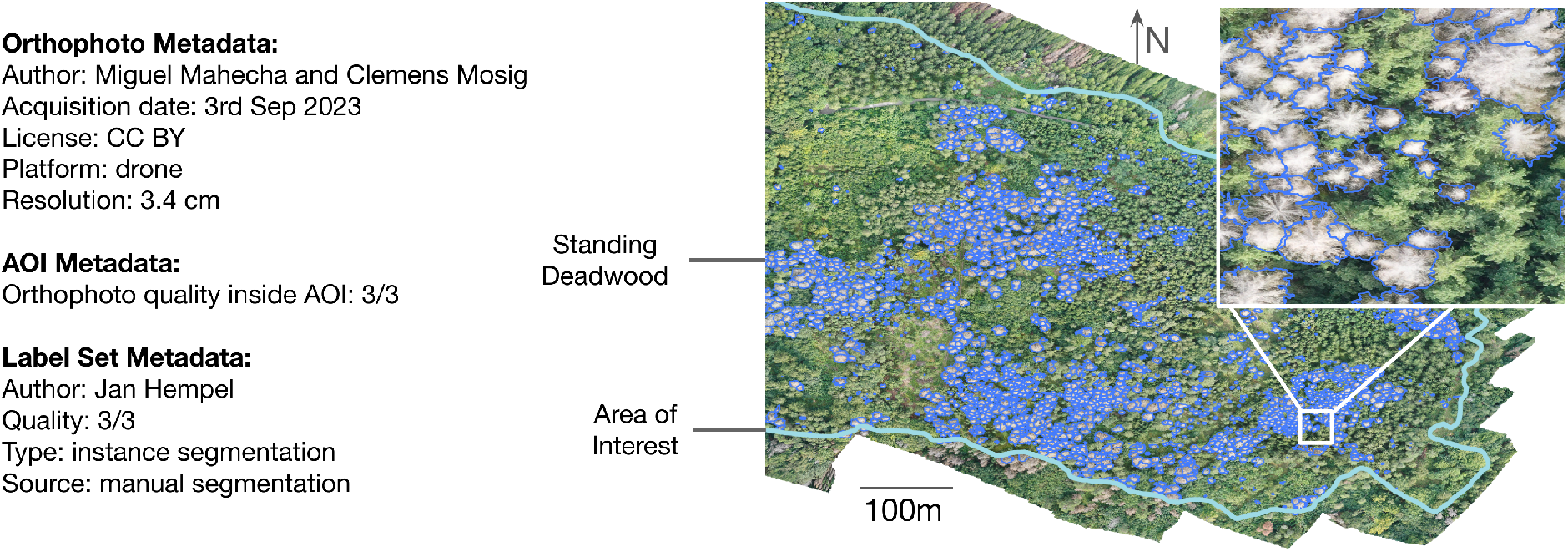
Sample entry of [orthophoto](https://deadtrees.earth/dataset/2491) (Jena, Germany, centroid: 50.911271°N 11.509977°W) with one label set for one area of interest (AOI) in the deadtrees.earth database. Only a simplified set of attributes are shown, see Figure 8 for the precise database structure.

Regardless of the spatial resolution, the information quality of an orthophoto can be constrained by various factors. These constraints include poor lighting conditions (*e.g*., underexposure), reconstruction artifacts, motion blur, or data gaps (Dandois et al. 2015; Frey et al. 2018). The image condition can vary heavily across an orthophoto, *e.g*., image edges are often distorted. To account for this, we assign each orthophoto an **area of interest (AOI)** that is a multi-polygon. This AOI object includes a score noting the quality of the orthophoto inside the AOI (see Figure 4). The scoring system ranges from 1 to 3, with 3 indicating near-perfect image quality, where only small portions (up to 5%) of the image are affected by constraints. A score of 2 is given if up to 25% of the AOI is affected, while a score of 1 is assigned when up to 50% of the orthoimage inside the AOI is constrained. Both the AOI and quality score are determined during a meticulous manual audit.

**Label sets** are polygons or points located over standing deadwood in orthophotos identified through visual inspection or from automatic segmentation (Cheng et al. 2024; Junttila et al. 2024; Schiefer et al. 2023). More specifically, there are four types of labels: (*i*) centroids of individual dead tree crowns, (*ii*) bounding boxes of individual dead trees, (*iii*) delineations of individual dead tree crowns (instance segmentation), and (*iv*) delineations around a group of adjacent dead trees or dead tree parts (semantic segmentation). Each label set is associated with an AOI, that also acts as boundary of the labeling effort. This means area inside the AOI that was not marked as deadwood can be assumed to be alive or non-tree objects (see Figure 4). Lastly, there can be multiple sets of labels from different sources for the same orthophoto, *e.g*., one may have been created manually while a second set was machine-generated by a segmentation model.

The quality of the labels will be assessed during an audit, where, again, a quality score between 1 and 3 will be assigned. A score of 3*/*3 means accurately delineated standing deadwood and partial dieback (see Figure 4). In the score of 2*/*3 we include sets where the vast majority of deadwood is labeled and/or delineations have imperfections, *e.g*., partially include forest floor or disregard partial dieback. Label sets with a score 1*/*3 include all other sets and are recommended to be excluded in further analysis or machine learning applications.

### 2.3 Platform architecture

The deadtrees.earth platform is an integrated web-based system designed to facilitate visualization, participation, management, and access to the deadtrees.earth database. The platform architecture con-sists of the following components: a user-facing front-end application, a cloud-hosted database for metadata and labels, a storage server for orthophotos and Cloud Optimized GeoTIFFs (COGs), a processing server for generating COGs, and user authentication (see Figure 5).

**Figure 5.**
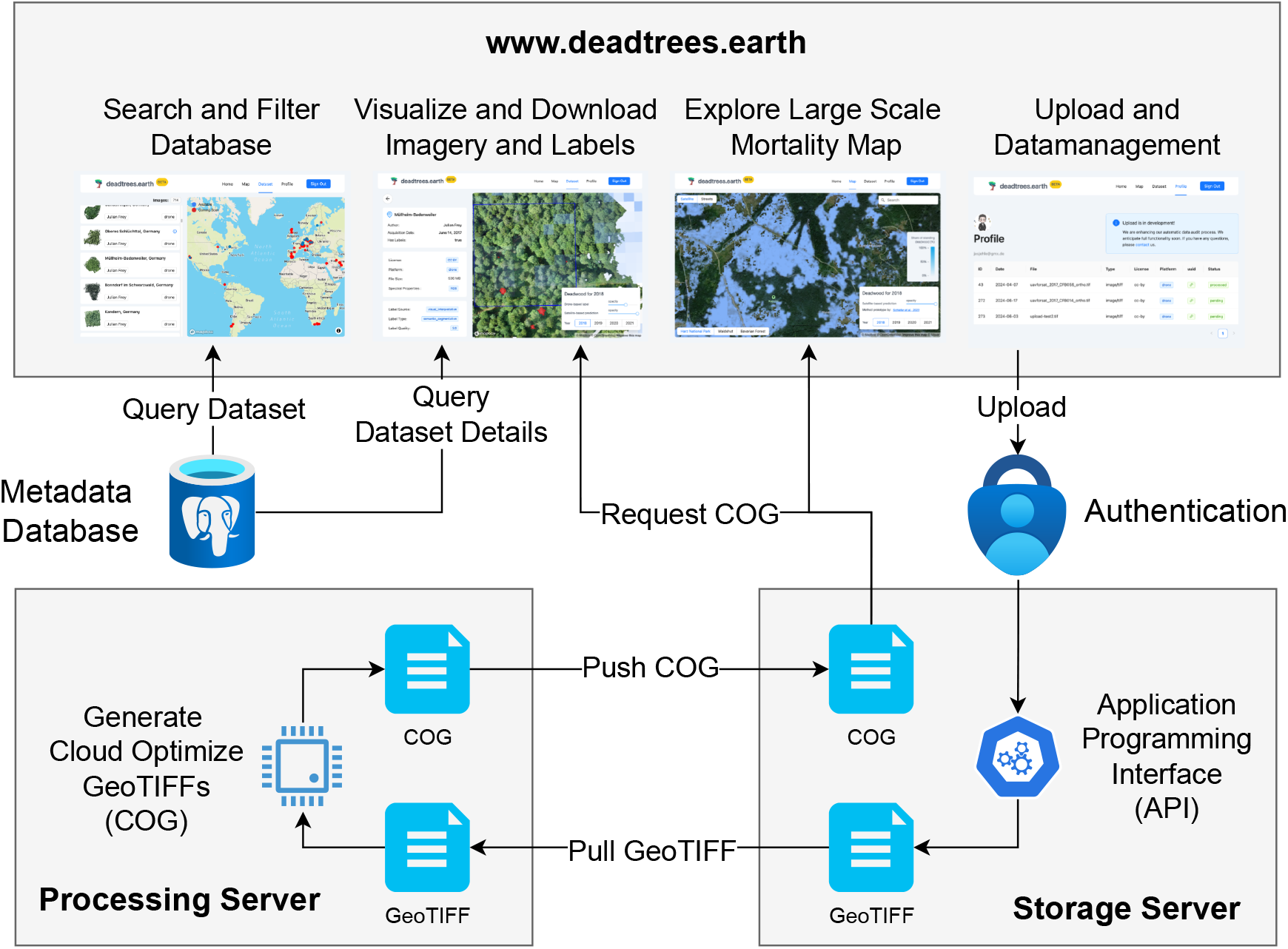
System diagram illustrating the main components of the deadtrees.earth platform and their interactions. Users can search and filter the database, visualize and download orthophotos, and explore a large-scale mortality map. The processing server generates Cloud Optimized GeoTIFFs (COGs) by pulling GeoTIFF files and pushing processed COGs to the storage server.

The front-end of the platform includes a landing page introducing users to the platform’s features, and a dataset page for searching and filtering the database through a list or world map. Users can select a specific dataset to access the *details page*, which visualizes one orthophoto with corresponding labels and their metadata. From here, users can download datasets without needing an account. A second page visualizes large-scale satellite-based deadwood maps. Finally, a user-specific profile page, which requires login, enables users to upload orthophotos and labels and manage their data.

Registered users can upload orthophotos, in the form of GeoTiffs, and labels to the system together with a set of metadata data that includes the author names and acquisition date per orthophoto. Upon successfull submission to the system, additional metadata is generated, that is administrative level, file size, file type. All metadata, along with vector labels, is stored in a cloud-hosted [Supabase](https://supabase.com/) database, which is accessible via Python and JavaScript client libraries. Data audit workflows require specific user access levels, which are assigned to the deadtrees.earth core team. For user authentication, we use Supabase Auth, which is based on JSON Web Tokens (JWTs). This ensures secure access while integrating with Supabase’s database features to implement Row Level Security (RLS), ensuring that each user can only access data they are authorized to view.

To efficiently visualize a large collection of orthophotos with minimal resources, the platform uses Cloud Optimized GeoTIFFs (COGs). COGs allow users to view and work with large orthophotos quickly and efficiently, which is especially helpful when bandwidth or processing power is limited. COGs are internally tiled and include overviews, making them accessible via HTTP range requests without the need for server-side processing. This approach allows clients to fetch only the necessary data, optimizing transfer and reducing server load. As a result, COGs significantly improve performance compared to traditional Web Map Services (WMS) such as GeoServer or MapServer.

The resource-intensive generation of COGs is performed on a separate processing server. The server periodically pulls user-uploaded GeoTIFF files from the storage server, performs the necessary processing, and pushes the generated COGs back to the storage server (see Figure 5). A Python-based REST API built with FastAPI manages processing tasks, user management, and resource allocation. The front-end initiates tasks such as uploading, downloading, metadata generation, and processing COGs through this REST API, which can also be used directly for programmatic data ingestion and processing. The deadtrees.earth API also employs a queuing system to manage processes and prevent downtime which ensures stability and scalability.

Finally, the platform’s modular design allows for future integration of advanced workflows, such as machine learning models for automated deadwood segmentation from drone imagery. By leveraging powerful local processing servers, these workflows can be added seamlessly, making the platform adaptable and flexible to meet evolving needs.

### 2.4 Data Sources and Current State of the Database

The primary sources for the orthophotos and labels are community contributions, *i.e*., datasets that individuals or institutions actively contributed. Given the large interest in monitoring tree mortality dynamics worldwide, the deadtrees.earth database received tremendous support from a wide array of individuals and institutions. So far, 87 institutions shared data across 67 countries.

#### Crowd-Sourcing

In addition to community contributions, the database integrates crowd-sourced data, *i.e*., datasets already freely available online. Indeed despite extensive community efforts to date, significant portions of the Earth remain uncovered in our database. Therefore to maximize database coverage, we integrate publicly available databases that adhere to appropriate licensing schemes.

While other initiatives, such as [GeoNadir](https://geonadir.com/), OpenAerialMap, and [OpenDroneMap](https://www.opendronemap.org/), also collect drone orthophotos, only OpenAerialMap currently ensures that all contributions are licensed under CC BY, making them suitable for use in projects like deadtrees.earth. As of June 2024, OpenAerialMap hosts over 15,000 aerial orthophotos. We use this community-driven resource to expand the deadtrees.earth database. However, most of the contributions to OpenAerialMap do not meet our database criteria due to limitations in resolution, site relevance, quality, or acquisition timing. To be able to extract usable images, we downloaded a summary of the metadata on 24th April 2024 through their open API. Then we first filter the entries with where at least 30% is covered by forest according to ESA Worldcover (Zanaga et al. 2022). To then remove orthophotos that lack the necessary spatial resolution (Figure 1), we filtered images to include only resolutions better than 10 cm, yielding 1102 samples. To only include orthophotos of forests within the growing season, we filtered the months May to August for samples north of latitude 23.5°N, December to March for samples south of latitude 23.5°S, and included all images for latitudes in between. Note that at a later stage we will differentiate between wet and dry seasons for tropical region. Finally, we manually iterated through the thumbnails or the original GeoTIFF of every orthophoto to visually check their quality. This resulted in a final set of 448 (out of *>* 15, 000 on OpenAerialMap) orthophotos with wide temporal (2007 to 2024) and geographic coverage (see Figure 6).

**Figure 6.**
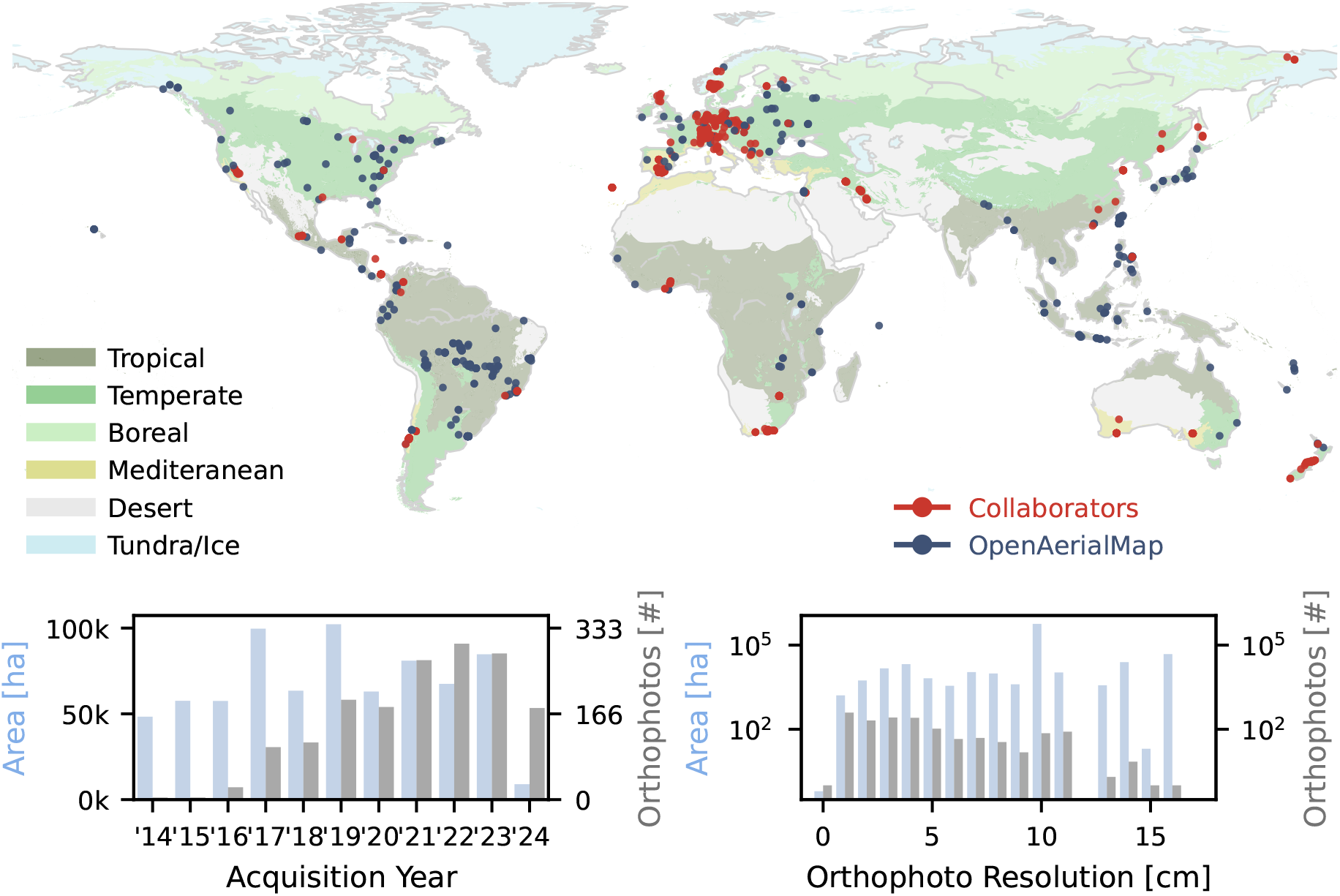
Initial statistics of the database upon launch depicting geographical, temporal, and resolution diversity. In the two bottom panels, drone orthophotos are accumulated by area (light blue) and count (dark gray). Different colors in the background depict different biomes (Olson et al. 2001).

**Figure 7.**
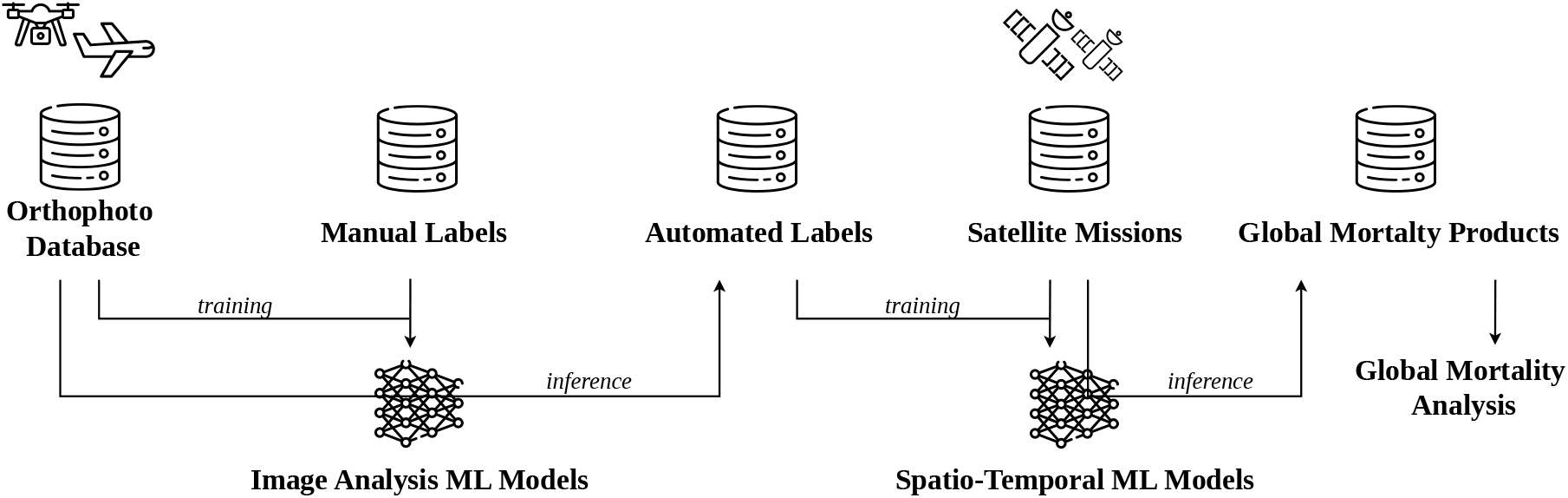
Generalized workflow to derive a global tree mortality product through the deadtrees.earth database.

It is worth noting that the dataset extracted from OpenAerialMap has a bias towards forests near human settlements, potentially over-representing ecosystems that might not be representative of the region. For example, an orthophoto may contain 20 ha of a relevant forest, but another 100 ha of the image contains a building site that the drone operator originally planned to capture. Nevertheless, this crowd-sourced dataset provides valuable, high-resolution imagery of forests in ecosystems that would otherwise not be part of our database. Additionally, this bias may provide an opportunity for studies focusing on studying forest fragments and urban forests. As OpenAerialMap grows in the future, we will continuously monitor their database for relevant submissions. Also, other relevant sources with a CC-BY license will be integrated.

#### Database Statistics

We launch the seed database with 1,390 centimeter-scale orthophotos covering 345,595 ha and spanning all continents (except Antarctica) through community contributions and crowd-sourced data. By the time of submission (Oct. 2024), the database consists of 998 (71%) drone orthophotos from community contributions and 392 (28%) crowd-sourced orthophotos extracted from OpenAerialMap (Figure 6). The increasing ease of use of drones within the last decade is reflected in the greater number of unique orthophotos in recent years. Additionally, the database includes 140 aerial images with resolutions less than 10 cm (Figure 6). Beyond local forest plots, we provide access to aerial images with machine-learning generated tree mortality labels that were published on our platform as the result of several studies (Cheng et al. 2024; Schwarz et al. 2024; Weinstein et al. 2024). These products cover the state of California (USA), Luxembourg, and 23 [NEON](https://data.neonscience.org/) sites in the USA (not shown in Figure 6).

#### Notable Collections

Although a large part of the database consists of individual locations that have been captured, it also features noteworthy collections that provide independent value, for example through temporal coverage across multiple months or years. Notable collections include:

- **Barro Colorado Island (Panama)** 90 orthophotos capturing the same 50 ha plot across 6 years (Vasquez et al. 2023).
- **Quebec (Canada)** Seven consecutive orthophotos of the same lake area from May to October 2021 (Cloutier et al. 2023, September).
- **Nationalpark Black Forest (Germany)** A 10-year timeseries covering the entire national park (Christoph Dreiser).
- **Baden Wuerttemberg (Germany)** 135 unique plots (*>* 1 ha) in southwest Germany captured in up to three different years, respectively ([ConFoBi](https://confobi.uni-freiburg.de/de/projekte)).
- **Andalucia (Spain)** 60 tree mortality sites (*>*15 ha) in otherwise protected national parks in 2023 (Clemens Mosig and Oscar Pérez-Priego).
- **Eastern Cape (South Africa)** 35 tree mortality sites captured between 2022 and 2024 providing unique data from Africa (Alastair Potts).
- **Zagros Forests (Iran)** 16 RGB Orthophotos captured in ca. 1 ha sample plots representing Quercus brantii (oak) decline. Distributed over the large latitudinal gradient of semiarid Zagros Forests in western Iran (Ghasemi et al. 2022, 2024a,b).
- **NIBIO UAV archive (Norway)**: 50 UAV RGB orthophotos captured by NIBIO’s Forest and Forest Resource division between 2017 - 2022 using a variety of DJI drones. These data were in collected primarily in south eastern Norway (Bhatnagar et al. 2022; Puliti et al. 2019, 2020).

The latter six collections have not been available to the public until now.

#### Labels

The database contains 54,320 *manually delineated* polygons delineating partial dieback, individual trees or multiple dead tree crowns. In total, 493 orthophotos and 58,219 ha are fully labeled, of which 245 have quality *3/3*, 231 have quality *2/3*, and 5 orthophotos have quality *1/3* (see Subsection 2.2 for quality definition). These datasets will soon be available as machine learning ready datasets (see Section Section 3) to support the community with training semantic or instance segmentation models. At present, this unique data collection would result in more than 600.000 labeled 512×512 patches or 170.000 labeled 1024×1024 patches.

For this data collection we strictly adhere to the FAIR principle (Wilkinson et al. 2016). All data is **F**indable, *i.e*., has a unique identifier, is described with metadata, and thus searchable. **A**ccess is provided through industry-standard and authentication-free HTTP requests on the website or programmatically (compare Subsection 2.3). We provide data **I**nteroperability by using GeoTIFF format and standard datatypes for metadata (see Figure 8). Lastly, all data is **R**eusable as it is published under a Creative Commons license.

**Figure 8.**
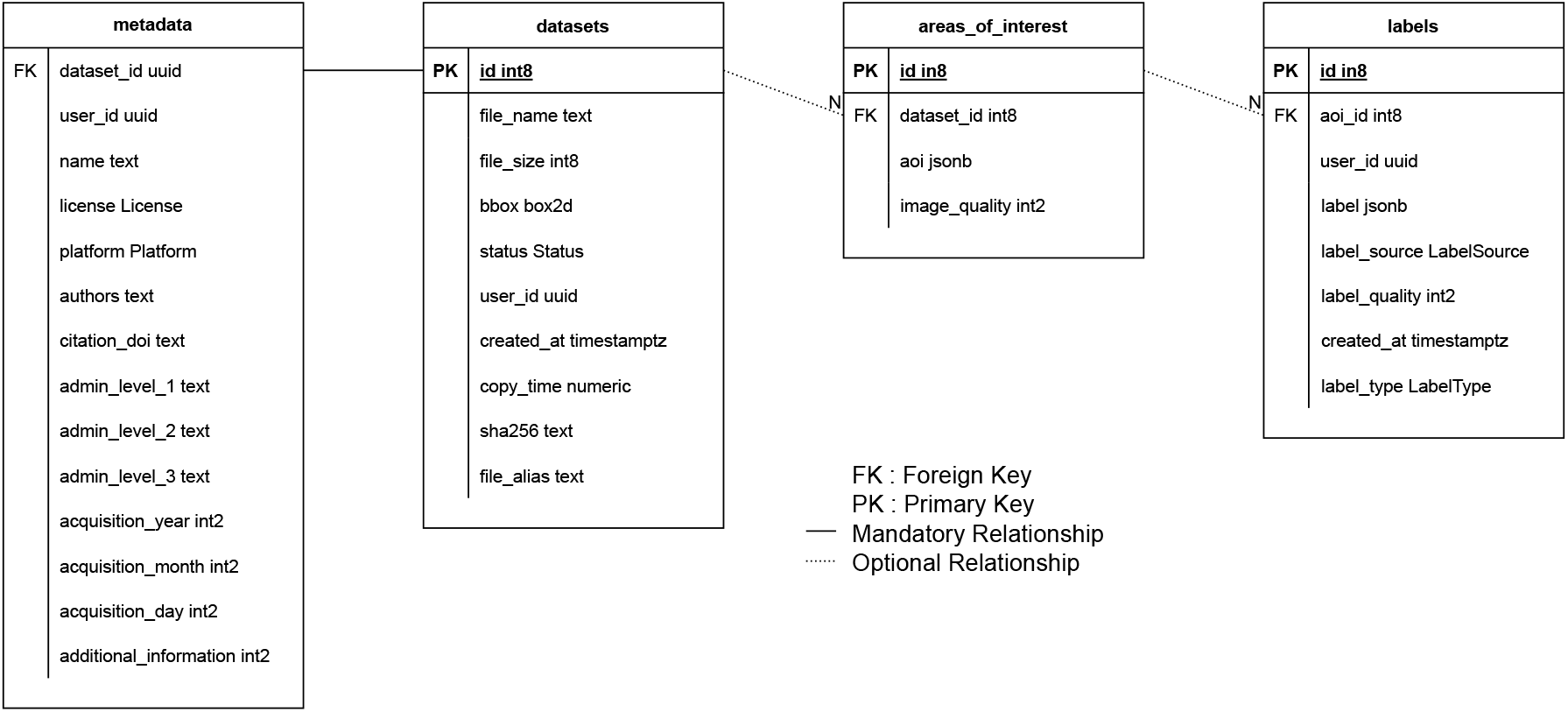
Full relational diagram of the deadtrees.earth database.

In summary, through community efforts and crowd-sourcing of data, and to the best of our knowledge, the deadtrees.earth database curates an unprecedented amount of super-resolution optical imagery and corresponding labels. With the increasing recognition of this database and the general growing willingness for open data in science and the public, we expect this database to continue expanding rapidly.

## 3 Outlook and Perspective

### 3.1 Database Expansion Through Community Contribution

Excess tree mortality is a global phenomenon whose underlying complexity can only be effectively assessed through community effort (The International Tree Mortality Network et al. 2024). The deadtrees.earth platform initiates with a collection of centimeter-scale forest orthophotos that is already orders of magnitude larger in spatial coverage and diversity than in any mortality-related study used. However, this collection is biased towards the Global North, and regions in Asia and Africa are particularly underrepresented (see Figure 6). As we aim to grow into a representative collection of tree mortality in the World’s forest ecosystems, we require a more diverse collection of orthophotos. We therefore encourage everyone in every community to take the opportunity to participate in this global initiative.

In the primary use case, a contributor submits an orthophoto covering any forest with a resolution better than 10 cm. Optionally, delineated standing deadwood can be submitted as shapefiles or similar formats. Beyond that, we also welcome lower-resolution aerial images with already delineated standing deadwood. These delineations can be manually obtained or also the product of automated segmentation, and need to be declared as such, *e.g*., the results of Cheng et al. 2024 are available in the database. The orthophotos do not necessarily need to contain large or any fractions of standing deadwood, as the machine learning models have to be trained on alive and dead trees. Since anyone can submit data to the database, a database manager manually reviews the supplied metadata and the geolocation of the orthophoto and, if available, grades the quality of the submitted label set. This ensures that the database continues to grow without barriers while maintaining the highest possible quality.

Newly submitted orthophotos of local tree mortality events bolster the global and temporal representativeness of the database. This is critical for training models that aim for a global transferability (Kattenborn et al. 2022; Meyer and Pebesma 2022), be it computer vision models that segment dead trees in drone data or satellite-based models. Hence, an individual submission of a user’s local forest can be an important missing puzzle piece in creating a representative training dataset. Subsequently, machine-learning models will improve in the user’s local region, providing a strong incentive to contribute their data as they indirectly benefit.

### 3.2 Towards Tree Mortality Models and Products from Local to Global Scale

Delineated standing deadwood identified from large amounts of centimeter-scale orthophotos is a powerful data source for creating high-precision training data. Deadtrees.earth provides a unique dataset that will enable the machine-learning community to create models and maps that are transferable at a global scale and robust across the diversity of forest ecosystems (Figure 7).

Given the rich database presented here, users can train various types of **computer vision models for identifying standing deadwood in drone orthoimagery**, *e.g*., in the form of semantic segmentation (polygons of dead crowns, twigs or branches), object detection (bounding boxes of individual trees), or instance segmentation (precise crowns of individual trees). With such models, one can perform inference on all orthophotos in the database to automatically reveal the local distributions of standing deadwood. This is particularly relevant for orthophotos that do not have labels from a human interpreter. Machine-learning-based predictions may even be advantageous over labels from human interpreters as they might be more standardized and objective (in contrast to manually delineated polygons from different human interpreters). This automated mapping of standing deadwood is also meant to be one of the core incentives for users interacting with the deadtrees.earth. Thus, deadtrees.earth will provide a hub for making machine-learning-based technology developed by the community accessible for non-experts (*e.g*., practitioners, citizens, Non-government organizations) or people with limited resources.

The local patterns of standing deadwood derived from orthophotos can be used as a reference for **large-scale machine-learning-based mapping using satellite data** from Sentinel, Landsat, or future satellite missions. While Landsat and Sentinel data are much coarser in resolution than drone data, approximately 10 m to 30 m, respectively, they have the advantage of having global coverage and being multi-spectral data. The temporal continuity of Sentinel or Landsat data supports the creation of accurate global products, as machine-learning models can harness the temporal and spectral patterns. For example, in optical satellite imagery, standing deadwood may look visually similar to a grayish forest floor or rocks (Figure 1). However, in a time series of multiple years, a dead tree can be differentiated from a forest floor or rocks based on its spectral history (Schiefer et al. 2023). This way, deadtrees.earth will provide satellite-based models and predictions at a global scale in the future.

To stimulate the development of machine-learning models for analyzing drone and satellite data, deadtrees.earth will provide ML-ready datasets, *e.g*., integrated into the [torchgeo](https://github.com/microsoft/torchgeo) library (Stewart et al. 2022). This will enable the community to develop and benchmark different methods effectively. Incentives for this might be further propelled by related coding competitions. Moreover, the machinelearning-ready datasets will enable the development of workflows that are directly compatible with the deadtrees.earth ecosystem, so that models and workflows developed in the community can be directly integrated as an application.

With the launch of deadtrees.earth we aim to attract a variety of communities to this multifaceted platform. Through simple, interactive visualizations of orthophotos together with labels and satellite-derived products on the website, we truly enable anyone to explore our and others’ tree mortality-related products. Viewing centimeter-scale imagery and satellite products side-by-side will enable benchmarks, validation, and finally an understanding of large-scale patterns of forest mortality. In a citizen science approach, non-specialists can also contribute data without prior knowledge of machine-learning methods used for further processing by us and the broad scientific community. In the future, we aim to further increase participation on deadtrees.earth by enabling users to delineate standing deadwood manually, correct AI segmentation outputs, and flag faulty predictions in the satellite data.

### 3.3 Applications of Global Tree Mortality Products

Global, high-quality tree mortality products can be used with environmental layers to attribute mortality dynamics to respective drivers and understand the variation in tree mortality dynamics. The variety of global tree mortality products that can be derived from the database will be a key component in enabling researchers to answer pressing questions: *Why are trees dying in the first place and how do the drivers (co)vary across tree species, ecosystems, or biomes? Why do some areas experience excess tree mortality while similar areas experience greening? Is tree mortality dependent upon the species or diversity of neighboring trees? What is the anthropogenic contribution to excess tree mortality? How long does standing deadwood remain in different ecosystems and does this relate to large-scale carbon balances? Where can tree mortality be attributed to global warming and climate extremes? Do the latter factors facilitate (invasive) pests and pathogens?* Given high product quality and increasing global coverage, we hope to support research on tree mortality from a local to a global scale and across biomes.

For example, one can combine standing deadwood maps with large-scale biomass maps (Santoro et al. 2020; Shendryk 2022) to facilitate our understanding of carbon fluxes. Given the temporal dynamics of standing deadwood, we can compare results to the outputs of vegetation models (*e.g*., (Köhler and Huth 1998)). Thereby, using remote sensing derived products to evaluate and also fine-tune or initialize parameterizations of vegetation models. Beyond Now- and Hindcasting, Forecasting of tree mortality should be possible if the community finds effective environmental predictors such that tree mortality dynamics for the subsequent year can be modeled.

Beyond tree mortality applications, we envision the orthophoto database to be used in a variety of other use cases. Since in general, this is a centimeter-scale orthophoto database of forests, one can also attempt to detect tree species, analyze tree line patterns, derive tree/non-tree products, pioneer studies on tree health, tree phenology, or attempt to track forest cover dynamics. Broadly speaking the general workflow (see Figure 7) of upscaling to global products can also be attempted for the same use cases. Especially suited may be forest cover products, tree species distribution maps, or revealing tree loss by forest management or windthrows.

## 4 Conclusions

The deadtrees.earth database is a centimeter-scale orthophoto collection with standing deadwood delineations. Already, it comprises 1,390 centimeter-scale orthophotos with more than 55,000 deadwood labels from the last decade distributed across the entire globe. The dataset has unprecedented coverage, and through machine learning methods and global remote sensing satellite missions, the scientific community can leverage this dataset to create models and global datasets, unlocking the potential to effectively track tree mortality dynamics. Ultimately, these data in concert with environmental layers will enable the scientific community to answer pressing questions on tree mortality. To reach this goal, the platform www.deadtrees.earth encompasses an interactive online system that aims to exploit aerial and satellite imagery for uncovering spatial and temporal patterns of tree mortality at a global scale. The web platform supports and encourages uploading and downloading user-generated orthophotos optionally together with labeled standing deadwood. The vision of this platform is an improved understanding of tree mortality patterns and processes from local to global scales. And this vision can only be accomplished through the collective effort of citizens and researchers. The dynamic nature of this database is meant to continuously increase our capacity to detect and understand tree mortality patterns. We hope that through the services of deadtrees.earth, we can attract ample data input from geographic regions that are currently still underrepresented (*e.g*., the global south). Finally, with this initiative, we support the paradigm shift in data-sharing practices in the scientific community.

## Acknowledgements

The study has been funded by the German Aerospace Centre (DLR) on behalf of the Federal Ministry for Economic Affairs and Climate Action (BMWK) under the projects *UAVforSAT* (project no. 50EE1909A) and *ML4Earth* (FKZ 50EE2201B). Further funding was received from the German Research Foundation (DFG) under the project *BigPlantSens* (project no. 444524904) and PANOPS (project no. 504978936). Further funding was received from the Ministry of Food, Rural Areas and Consumer Protection under the project PRIMA (project no. 52-8670.00). Some of the icons were provided by Flaticon. JF acknowledges funding by the German Research Foundation (DFG Project ConFobi, GRK 2123). CM, MDM, and JU acknowledge the financial support by the Federal Ministry of Education and Research of Germany and by Sächsische Staatsministerium für Wissenschaft,Kultur und Tourismus in the programme Center of Excellence for AI-research, Center for Scalable Data Analytics and Artificial Intelligence Dresden/Leipzig, project identification number: ScaDS.AI. CM and MDM thank the European Space Agency for funding the “DeepFeatures” project via the AI4SCIENCE activity. SH and YC are funded by Villum Fonden (DRYTIP project, grant agreement no. 37465) and the University of Copenhagen (PerformLCA project, UCPH Strategic plan 2023 Data+ Pool). We acknowledge the Black Forest National Park Administration as on of the data providers. The research of KCC was carried out at Oak Ridge National Laboratory, which is managed by the University of Tennessee-Battelle, LLC, under contract DE-AC05-00OR22725 with the U.S. Department of Energy. This study was supported by the International Tree Mortality Network (https://tree-mortality.net/).

## Supplementary Material

## Author contributions

Conceptualization: C. Mosig, T. Kattenborn, and J. Vajna-Jehle. Writing - original draft: C. Mosig and T. Kattenborn. Writing - review & editing: C. Mosig, Y. Cheng, J. Vajna-Jehle, S. Junttila, T. Kattenborn, H. Hartmann, S. Horion, M. D. Mahecha, D. Montero, M. B. Schwenke. Analysis and Visualization: C. Mosig, T. Kattenborn, and J. Vajna-Jehle. Platform development (front-end, back-end): J. Vajna-Jehle, M. Mälicke, C. Mosig. All others contributed data, revised the manuscript, and gave approval for publication.

